# Protocol for Single-Cell Patch-Proteomics: identify specific proteins associated with electrophysiological properties from individual neurons

**DOI:** 10.64898/2026.09.24.750455

**Authors:** J. Christian Althaus, Yiding Cao, Yang Song, Ali Haghani, Rachel Callaghan, Shannon J. Moore, Sarah J. Parker, Catherine C. Kaczorowski

## Abstract

Single-cell patch-proteomics is a novel and powerful technique combining electrophysiological and quantitative proteomic characterization from individually sampled neurons. Following whole-cell patch clamp characterization of neuronal excitability, we collect intracellular contents and, using the latest advancements in Liquid Chromatography and Mass Spectrometry, identify and quantify peptides and their corresponding proteins. By cell-matched one-to-one associations between electrophysiological properties and protein abundance, this approach provides a means to elucidate molecular mechanisms of neuronal function on individual neuron basis.

**Graphical Abstract:** 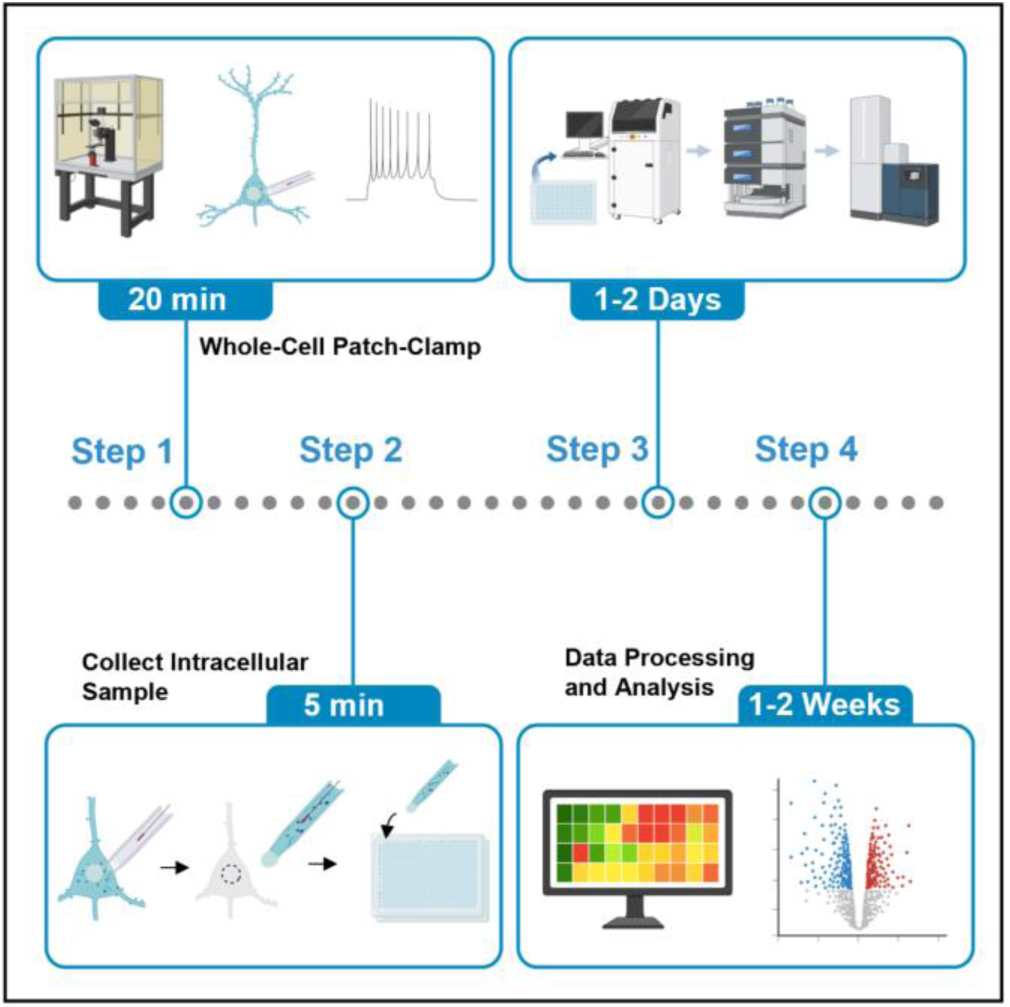

## Innovation

Neuronal populations exhibit heterogeneity in form and function, even within the same brain region. Variation in spiking properties, intrinsic excitability, plasticity, and other neuronal features arise from distinct expression of ion channels, receptors, and other proteins that are dynamically regulated and allow neural networks to optimize computations for enhanced learning and memory (Dunn et al., 2018; Graves et al., 2016; Jarsky et al., 2008; Josselyn & Tonegawa, 2020; Kaczorowski et al., 2012; Kaczorowski & Disterhoft, 2009; Walker & Frost, 2024). This complexity highlights the necessity to precisely identify and understand how proteomic signatures of individual neurons within neural circuits mediate learning and memory in healthy and diseased brain states.

The predecessor to patch-proteomics is the patch-sequencing technique, which measures RNA transcript along with electrophysiological and/or morphological properties from a single neuron. Because RNA can be amplified, it is relatively straightforward to generate transcriptomic profiles from minute single-neuron samples, which has enhanced the specificity and granularity of single-cell analysis. However, while it was initially thought that protein expression changes could be inferred based on transcriptomic data (Cadwell et al., 2016, 2017; Lee et al., 2021; Lipovsek et al., 2020), it has become clear that the correlation of mRNA and protein abundance is often poor – especially in the context of brain aging and neurodegenerative diseases (Pascal et al., 2008; Schimanski & Barnes, 2010; Schwanhäusser et al., 2011; Tasaki et al., 2022; Upadhya & Ryan, 2022; Vogel & Marcotte, 2012). This is due, in part, to post-translational modifications, protein degradation via the ubiquitin proteasome system, protein folding, and targeted localization that can result in a dramatically different proteomic profile relative to genomic and/or transcriptomic features (Deribe et al., 2010; Hartl et al., 2011; Liu et al., 2016; Nakai et al., 1999; Wang et al., 2013). It is therefore crucial to interrogate protein abundance directly and not rely on RNA as a proxy – despite what the central dogma may suggest – to be able to link changes in protein expression with the functional state of the neuron.

Until recently, proteomic analysis from individual neurons had remained elusive. This is in large part because protein, unlike DNA and RNA, cannot be amplified through molecular biology techniques, making detection of proteins from miniscule single-neuron samples extremely difficult. Recent advancements in the field of Liquid Chromatography and Mass Spectrometry (LC-MS) have now provided the means to measure these small amounts of protein from individual cells (Kreimer et al., 2023; Saddic et al., 2025), which has thus been termed “patch-proteomics”.

This protocol is specifically tailored towards users who have prior experience in patch-clamp electrophysiology in acute brain slices and want to acquire proteomic data from the same cells. We demonstrate this approach by measuring the abundance of specific proteins from individual neurons collected following whole-cell patch clamp electrophysiological characterization of intrinsic excitability, and associate expression levels of unique proteins with neuronal function. While beyond the scope of the protocol presented here, this approach could be further extended to correlate neuronal proteomic signatures to other parameters of interest, such as morphological properties (on a cell-by-cell basis) or behavioral performance (such as cognitive function, on a by-animal basis). Taken together, this powerful patch-proteomics approach lends itself to unbiased discovery and novel insights into molecular mechanisms involved in basal neuronal function as well as memory- or disease-related plasticity across a range of network states and computations.

For a step-by-step protocol on how to perform whole-cell patch clamp in acute brain slices, we recommend following the previously published STAR Protocol on this technique (Chen et al., 2025; doi: 10.1016/j.xpro.2025.104008). Below we briefly summarize these steps, noting both *“Non-essential differences”* and highlighting “C*ritical modifications*.” Non-essential differences provide context, but these minor variations are not expected to significantly affect the outcome. Critical modifications are crucial to the success of the protocol and should be followed exactly as written.

### Institutional permissions

This protocol and subsequent experiments were performed on UM-HET3 mice (RRID:IMSR_JAX:036603) at the University of Michigan in compliance with NIH guidelines and the University of Michigan Committee on Use and Care of Animals. Mice were on a 12h/12h light/dark cycle with ad libitum access to food and water. Performing this protocol requires following NIH guidelines and obtaining permissions from the relevant institutional committees for working with animal models.

### Solution preparation

Timing 1-2 days

#### Critical Modifications from Chen et al

1. Prepare lysis buffer

a. 100mM triethylammonium bicarbonate (TEAB), 0.2% n-dodecyl-β-D-maltoside (DDM), 50mM Tris pH 8.0
2. Load lysis buffer into loading vial of a Scienone CellenONE cell sorting and robotic liquid dispensing instrument
3. Dispense 200nL lysis buffer into each well of a 384-well plate
4. Spin plate at 500 relative centrifugal force (rcf) to ensure buffer reaches bottom of each well
5. Cover 384-well plate with cold storage foil and store at -80°C until sample collection begins

a. Store lysis buffer-filled plates no longer than 6 months at -80°C

Note: as an optional step, to further ensure minimal peptide adsorption, pre-coat plate wells with 0.2% DDM solution overnight, invert to remove excess coating solution and dry prior to lysis buffer dispensing.

#### Non-essential differences from Chen et al

1. Prepare potassium gluconate-based internal solution

a. 115 mM potassium-gluconate, 20 mM potassium chloride, 10 mM sodium phosphocreatine, 10 mM HEPES, 2 mM Mg-ATP, and 0.3 mM Na-GTP, pH adjusted to 7.3
2. Prepare the Recovery artificial Cerebral Spinal Fluid (aCSF)

a. 93mM NMDG, 2.5mM potassium chloride, 1.4mM Sodium phosphate monobasic, 30mM sodium bicarbonate, 20mM HEPES, 25mM D-Glucose, 3mM sodium L-ascorbate, 2mM thiouera, 6.1mM N-acetyl-L-cysteine, 3 mM sodium pyruvate, <0.1mM taurine, 10mM magnesium sulfate [heptahydate], 0.5mM calcium chloride [dihydate], adjust to a pH of 7.4, bubble in to saturate with 95% O_2_ / 5% CO_2_
3. Prepare the Holding aCSF

a. 94mM sodium chloride, 2.5mM potassium chloride, 1.4mM Sodium phosphate monobasic, 25mM sodium bicarbonate, 20mM HEPES, 25mM D-glucose, 3mM myoinositol, 5mM sodium L-ascorbate, 2mM thiouera, 6.1mM N-acetyl-L-cysteine, 3mM sodium pyruvate, <0.1mM taurine, 2mM magnesium sulfate [heptahydate], 2mM calcium chloride [dihydate], adjusted pH of 7.4, bubble in to saturate with 95% O_2_ / 5% CO_2_
4. Prepare the recording aCSF solution

a. 30mM sodium chloride, 0.625 mM potassium chloride, 0.625mM calcium chloride [dihydate], 0.25 mM magnesium sulfate [anhydrous], 0.3125mM sodium phosphate monobasic, 23mM sodium bicarbonate, 10mM D-glucose, adjusted pH of 7.4, bubble in to saturate with 95% O_2_ / 5% CO_2_

### Brain slice preparation

Timing 1.5-2 h

#### Non-essential differences to Chen et al

5. Anesthetize mouse using isoflurane.
6. No transcardial perfusion prior to decapitation and brain removal.
7. Acquire transverse hippocampal slices.

a. Separate hemispheres and block tissue on dorsal side of each hemisphere.
b. Glue hemispheres block side down on the slice plate so ventral side is cut first.
8. Incubate slices at 34^◦^C for 15 minutes in Recovery aCSF.
9. Transfer slices into Holding aCSF at room temperature for one hour.

### Patch-clamp rig setup

Timing 15 min

#### Critical modifications to Chen et al

10. Prepare the sample collection plate.

a. Fill a foam bucket ¾ full of pelleted dry ice.
b. Place the CoolRack XT 384 block on the dry ice with the wells facing up (Figure 1A).
c. Fill in extra pellets of dry ice around the CoolRack.
d. Wait a minimum of 20 minutes for the CoolRack to reach temperature.
e. Remove the lysis buffer-containing 384-well plate out of the -80°C freezer and immediately nest onto CoolRack (Figure 1B).
f. Cover foam bucket with lid and place on table next to electrophysiology rig.
11. Assemble the expulsion device.

a. Attach a syringe filter to the end of a 1mL syringe.
b. Connect 6-8 inches of TPE tubing to the other end of the syringe filter (a plastic adaptor may be required).
c. Attach the other end of the tubing to an electrode holder in which the electrode has been removed (Figure 1C).
d. Spray with ethanol, wipe down, and store in cardboard box when not in use.

**Figure 1.**
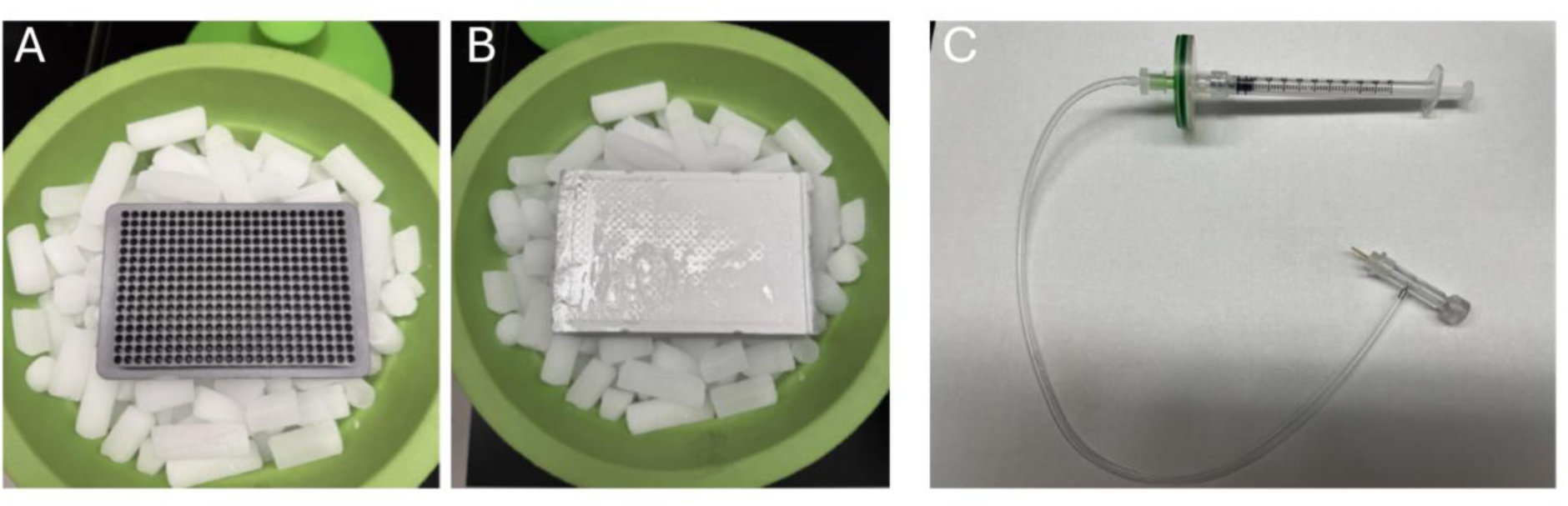
Setup for individual neuron protein sample collection. A) Placement of CoolRack over dry ice and allowed to cool for 20min. B) 384-well plate sealed with cold storage foil is taken out of -80°C freezer and nested onto CoolRack. C) Expulsion device: 1mL syringe, 0.88uM syringe filter, TPE tubing, and electrode holder.

Whole-Cell Patch-Clamping

Timing: 30 min

#### Critical modifications to Chen et al

12. Fill glass recording pipettes with internal solution
Use a P10 Eppendorf Pipettor with microloader pipettes and fill the glass pipette with 2.5uL of internal solution
Ensure all the solution is expelled into the beveled tip and remove any visible bubbles
Fill glass recording pipette with 2.5uL, ensuring all the solution is expelled into the beveled tip CRITICAL: When installing glass pipette onto the electrode, the end of the electrode must submerge in the small 2.5uL volume at the tip of the pipette. This may require altering the length of the electrode. It should be long enough to reach the internal solution, but not too long where it runs into the beveled end of the pipette, potentially breaking it. CRITICAL: During the recording period, keep an eye on the location of the recording electrode in relation to the soma. Move the pipette with small, fine-tuned adjustments throughout the recording process to maintain as close to the original patch position on the cell as possible. This will set up a successful extraction and retraction of the cytosol/nucleus. If the pipette drifts too far and stretches or punctures the cell too much, achieving a high-quality extraction will be difficult (See Troubleshooting 2).

## Key Resources Table

| REAGENT or RESOURCE | SOURCE | IDENTIFIER |
| --- | --- | --- |
| **Chemicals, peptides, and recombinant proteins** |  |  |
| Triethylammonium bicarbonate | Sigma-Aldrich | 18597 |
| DDM (n-dodecyl β-D-maltoside) (10%) | Invitrogen | BN2005 |
| Trypsin/Lys-C Mix | Promega | V5072 |
| Formic acid | Thermo Scientific | 28905 |
| Potassium D-gluconate (Potassium Gluconate) | Sigma-Aldrich | G4500 |
| Potassium Chloride (KCl) | Sigma-Aldrich | P9333 |
| Phosphocreatine disodium salt (Sodium Phosphocreatine) | Tocris | 4325 |
| HEPES | Sigma-Aldrich | H3375 |
| Adenosine 5'-triphosphate magnesium salt (Mg-ATP) | Sigma-Aldrich | A9187 |
| Guanosine 5'-triphosphate sodium salt hydrate (Na-GTP) | Sigma-Aldrich | G8877 |
| Water (Nuclease Free) | Fisher | BP2484 |
| Single-Cell Lysis Buffer | TaKaRa | 635013 |
| N-Methyl-D-glucamine (NMDG) | Sigma-Aldrich | M2004 |
| Sodium Phosphate monobasic | Sigma-Aldrich | S8282 |
| Sodium bicarbonate | Thermo Fisher Scientific | 424270010 |
| D-Glucose (anhydrous) | Fisher Scientific | D16 |
| Sodium L-ascorbate | Sigma-Aldrich | A4034 |
| Thiourea | Sigma-Aldrich | T7875 |
| N-Acetyl-L-cysteine | Sigma-Aldrich | A7250 |
| Sodium Pyruvate | Sigma-Aldrich | P2256 |
| Taurine | Sigma-Aldrich | T8691 |
| Magnesium sulfate (heptahydrate) | Millipore Sigma | M1880 |
| Calcium chloride (dihydrate) | Sigma-Aldrich | C7902 |
| Sodium Chloride (NaCl) | Sigma-Aldrich | S9888 |
| Magnesium sulfate (anhydrous) | Fisher Scientific | M65 |
| **Deposited data** |  |  |
| MassIVE Proteomics Data Repository Server | Center for Computational Mass Spectrometry, UCSD | https://massive.ucsd.edu/<br>MSV000102490 |
| **Experimental models: Organisms/strains** |  |  |
| Mouse: (CByB6F1/J x C3D2F1/J)/J (UM-HET3) | Jackson Laboratories | JAX: 036603 |
| **Software and algorithms** |  |  |
| DIA-NN v2.3.1 Academia | Demichev et al. | https://github.com/vdemichev/DiaNN/releases/tag/2.0 |
| FragPipe v23.0:<br>MSFragger v4.3<br>Percolator v3.7.1<br>Philosopher v5.1.1<br>EasyPQP v0.1.52 | Nesvilab proteome bioinformatics group | https://github.com/Nesvilab |
| Python v3.11.15 | Python Software Foundation | https://www.python.org/ |
| Scanpy v1.11.5 | scverse | https://scanpy.readthedocs.io/en/latest/ |
| AnnData v0.12.11 | scverse | https://anndata.readthedocs.io/en/latest/ |
| NumPy v2.4.4 | NumPy team | https://numpy.org/ |
| Pandas v2.3.3 | NumFOCUS | https://pandas.pydata.org/ |
| Statsmodels v0.14.6 | statsmodels-developers | https://www.statsmodels.org/stable/index.html |
| Matplotlib v3.10.9 | Matplotlib Development Team | https://matplotlib.org/ |
| Seaborn v3.10.9 | Seaborn Developer | https://seaborn.pydata.org/ |
| **Other** |  |  |
| Next Generation Micropipette Puller P-1000 | Sutter Instrument | P-1000 |
| This Wall Borosilicate Glass with Filament | Sutter Instrument | BF150-110-10 |
| Fisherbrand™ Sterile Syringes for Single Use, 1mL | Fisher Scientific | 14955464 |
| Sterile Syringe filter, 0.80uM | Corning | 431221 |
| 1-HL-U Electrode Holder Package | Molecular Devices | 1-HL-U |
| Hard-Shell® 384-Well PCR Plates, clear/clear | BIO-RAD | HSP3801 |
| Corning® CoolRack 96c0.5mL, Holds 96 x 0.5mL 2D Tubes | Corning | 432056 |
| Dry Ice | Cryogenic Gases | CYG1045 |
| Corning™ Round Ice Bucket with Lid, 2.5 L | Fisher Scientific | 07-210-134 |
| Microloader 2x96ST | Calibre Scientific | 930001007 |
| timsTOF Ultra 2 w/ Captive source (Mass Spectrometer) | Bruker | 1911342 |
| Column Toaster Max | Bruker | 1909136 |
| CaptiveSpray 2 Emitter 10um | Bruker | 1811112 |
| PepSep C18,<br>15cmx75µm,1.9µm (Analytical<br>column) | Bruker | 1893473 |
| Ultimate3000 LC:<br>WPS-3000TPL RS<br>(Autosampler)<br>NCS-3500RS CAP (Gradient<br>Pump)<br>HPG-3400RS (Loading pump)<br>SRD-3600 (Degasser) | Thermo Fisher<br>Scientific | 5826.0020<br>5041.0020<br>5040.0046<br>5035.9230 |
| EXP STEM TRAP, 0.17 µL,<br>PEP/10 µm (Trap column) | Optimize Technologies | 15-03992-TN |
| CellenONE version F1.4,<br>Software version 2.0 | Scienion | C00174 |
| Beckman ECHO ACCESS liquid<br>dispensing robot | Beckman Coulter Life<br>Sciences | 83493219 |
| PDC M Piezo Dispense Capillary<br>Uncoated | Scienion | C00016 |

### Step-by-step Instructions

#### Acquiring the cytosol and nucleus from a patched neuron

Timing 3-5 minutes

The goal of the following steps is to carefully remove the cytosol and nucleus from the cell with the glass micropipette. “Corking” the end of the pipette via a nucleated patch is important for ensuring the intracellular sample is uncontaminated by the surrounding tissue (For entire Extraction, Retraction process, see Video 1).

1. Extraction Phase

a. While in voltage clamp, run the membrane test in Clampex and note the membrane resistance (R_m_) and stability of the waveform.
b. If necessary, move the pipette in small, slow adjustments to the x, y, z center of the soma
c. If possible, visually locate the nucleus within the soma and keep an eye on it throughout the extraction process (Figure 2A). Note: throughout the recording process, it may become very difficult to visualize neuron morphology and nucleus. When in doubt, the best practice is to position the pipette in what appears to the x,y,z center of the soma prior to extraction.
d. Apply very light negative pressure to the syringe used for patching and maintain it by closing off the valve mid-suction (See Troubleshooting 1)
e. If membrane test waveform begins to destabilize, open the syringe valve, wait for stabilization, then reapply negative pressure as in step (d). Note: If the seal on the cell is completely lost (R_m_<50MΩ), we recommend continuing with the extraction, but making note for quality control (QC) purposes. Single-cell protein detection should still be possible, so long as the nucleus is extracted successfully.
f. Allow negative pressure to suction the cytosol for 10-20 seconds. The soma should begin to shrink slightly
g. If you can visibly locate the nucleus, move the pipette toward nucleus and push the tip up against the edge of the nuclear membrane
2. Retraction Phase

a. Move the pipette in the x-direction to the edge of the cell membrane (towards the manipulator side of the cell)
b. Switch the micromanipulator to “Approach” mode so the pipette will move in a diagonal direction (x and z). If “Approach” mode is not available, move the pipette step wise up and out of the soma
c. Retract the nucleus slowly through the cell membrane (Figure 2B.; See Troubleshooting 2)
  i. Retract 2-3 microns
  ii. Pause for 2-3 seconds to allow cell membrane to relax
  iii. Repeat steps i. and ii.
  1. i.e. Retract… Pause… Retract… Pause… Retract… etc. **CRITICAL:** Retraction of the nucleus should be done at a slow and deliberate pace with intervals of pausing to allow the cell membrane to “relax” and release the nucleus. Removing it too quickly or without pausing will result in the nucleus being ripped through the cell membrane or left behind in the soma
d. Once the nucleus is fully removed from the soma, a gradual increase in R_m_ occurs as a nucleated patch is formed and a 1GΩ seal is achieved (typically 10-20s to reach 1GΩ; Figure 2C.; See Troubleshooting 3)
e. Move efficiently in approach mode through and out of the tissue into the external aCSF solution just above the surface of the slice (Figure 2D) Note: Do not move too fast, the nucleus can become dislodged off the end of the pipette if it snags on debris, blood vessels, etc. **CRITICAL:** If retraction of the nucleus is unsuccessful, or it becomes dislodged or suctioned fully into the pipette while traversing the tissue, we recommend discarding the collected sample. “Corking” the end of the pipette with the nucleus is critical for single-cell interrogation. Suctioning extracellular tissue will result in a protein profile that is not isolated to the specifically interrogated cell and will confound any post-hoc analysis.
f. Apply stronger negative pressure and visualize the nucleus slowly being sucked into the pipette (Figure 2D)
g. Move the objective in the z-direction away from the tissue, but not all the way out of the bath Note: The surface tension ripple that occurs from removing the objective out of the bath can cause the nucleus to be physically dislodged from the pipette tip
h. In approach mode, carefully remove the pipette quickly from the bath, being sure to safely clear the objective and stage components
i. Fully remove the objective from the bath
j. Leave the pipette on the electrode hovering in the air while setting up for expulsion phase
3. Expulsion Phase

a. Ethanol gloves
b. Set the plunger on the expulsion device to 1.0mL
c. Peel back the foil on the 384-well plate to ensure wells can be accessed, but replace foil while retrieving pipette
d. Remove the pipette from electrode holder on micromanipulator and insert it into the expulsion device
e. While holding the pipette and expulsion device in one hand, remove the foil over the 384-well plate
f. Push down the syringe plunger on the expulsion device to apply 0.2mL of pressure (to 0.8mL) Note: the pressure applied during step (f) is not strong enough to expel the sample. Do this BEFORE placing pipette into well
g. Carefully, but deliberately, lower the pipette tip into the very bottom of the appropriate well and press the tip against the base off the well (Figure 3C). Note: the very tip of the pipette will remain in the well with no fluid left in the remaining pipette. Visually confirm the broken tip. The small volume of sample will freeze quickly upon contact with the inside of the well, so a swift break of the tip is imperative to ensure the sample expels or sticks to the inside of the well
h. Remove the pipette from well
i. Replace the foil on the well plate
j. Keep diligent sample records

i. Track which well-plate position corresponds to which electrophysiological recording files for each cell (and from each animal)
k. When sample collection for each day is finished, quickly wrap the open end of the 384-well plate in parafilm or aluminum foil to seal
l. Store plate in -80°C freezer until next use Note: We recommend using a single plate for 10-15 recording sessions/sample collections before starting a new plate. This is to avoid any potential freeze/thaw issues by continuously removing it from -80°C, even though the plate is on dry ice. Pause Point: store plates in -80°C freezer until next step. It is recommended to store them no longer than six months prior to running liquid chromatography and mass spectrometry.

**Figure 2.**
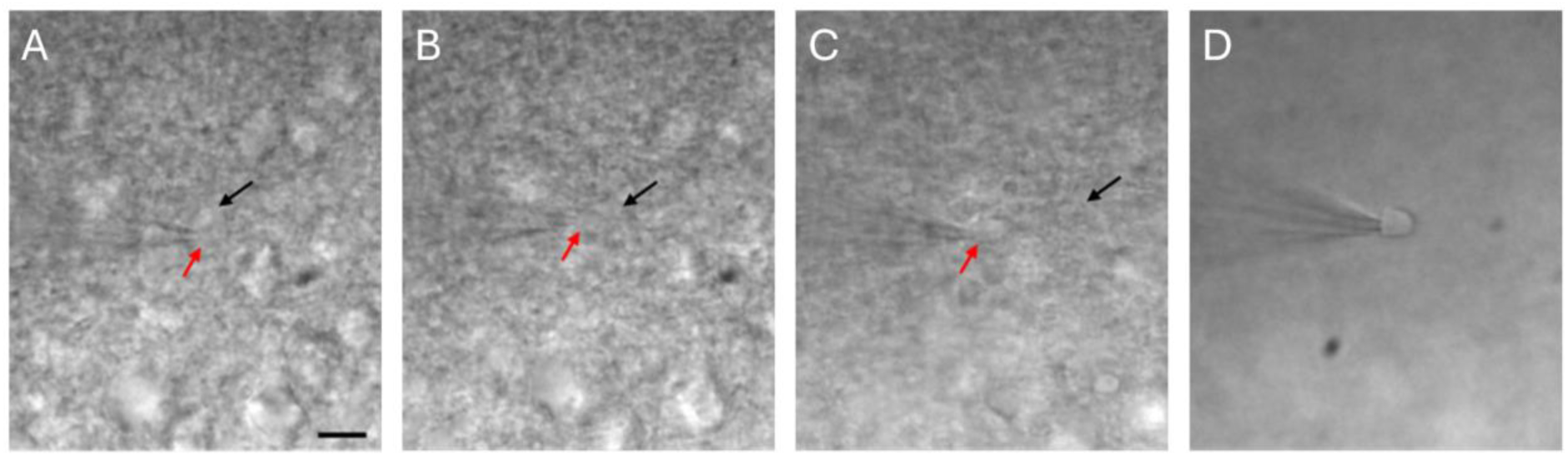
Extraction and retraction of cytosol and nucleus. A) Extraction. Visualize the nucleus (red arrow) inside the cell membrane (black arrow) and apply negative pressure to begin extraction. B) Retraction. The nucleus should follow the tip of the pipette as it begins to pull away from the cell membrane. C) Nucleated patch. As the nucleus is fully removed from the cell body a nucleated patch will form on the end of the pipette, sealing the intracellular sample inside. A high-quality extraction will result in a 1GΩ membrane resistance on the nucleus. D) Once fully retracted from the tissue, apply stronger negative pressure to ensure the sample adheres to pipette prior to removal from the bath (Scale bar = 10uM).

**Figure 3.**
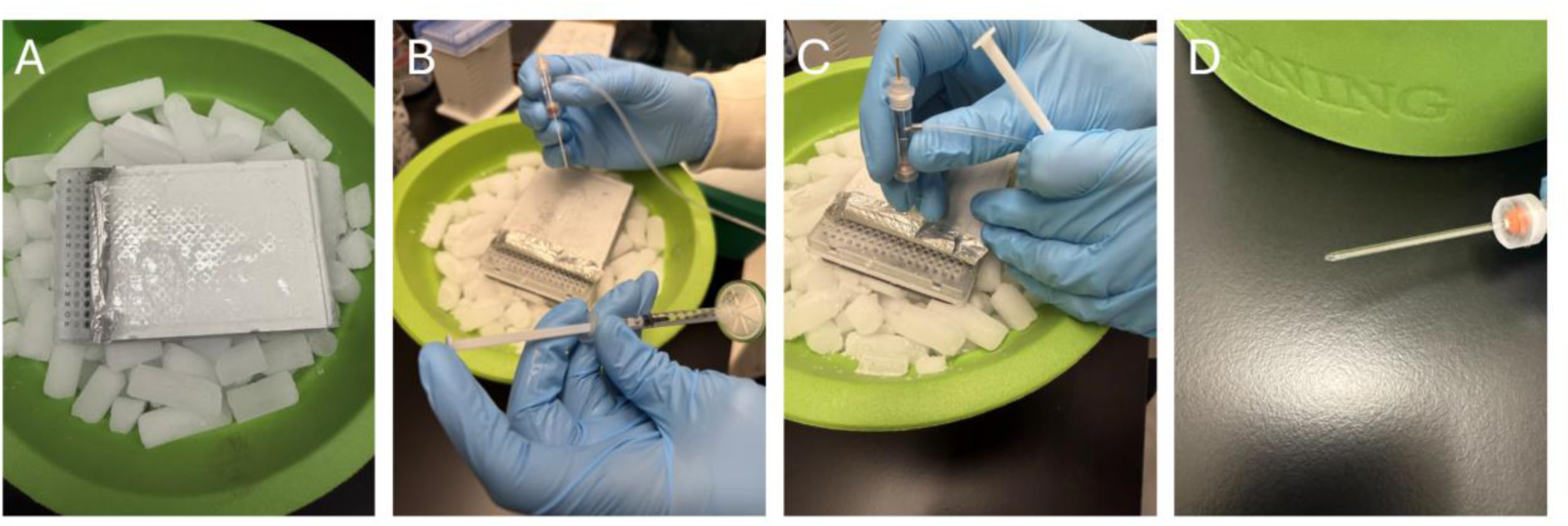
Expulsion of intracellular sample. A) Peel back the foil from the well plate. B) Insert the glass pipette containing the intracellular sample into the expulsion device and apply 0.2mL of positive pressure. C) Insert tip of pipette all the way to the bottom of the well and press firmly against the base of the well, breaking the pipette and leaving the tip with the sample in the bottom of the well with the frozen lysis buffer. D) Example of broken pipette following expulsion. Most of the beveled end should be left in the well.

### Detecting Single-Cell Proteins Using Liquid Chromatography Mass Spectrometry

#### Timing: 6 hours sample preparation per plate, 45 minutes data acquisition per cell

The capacity to detect single-cell proteomes requires highly sensitive LC-MS equipment. Recent advances in instrumentation have enable sufficient sensitivity to identify proteins directly in digested peptides from the lysates of individual cells (Bubis et al., 2025; Ctortecka et al., 2024). One of the most important innovations to minimize loss of peptides during sample processing is automated, nanovolume scale sample processing workflows with zero transfer or ‘clean up’ steps using MS-compatible buffers. These are designed to minimize adsorption to plastics in tubes and processing plates and maximize transfer of cellular proteins to the LC-MS system. These improvements to the workflow are key to successful single-neuron patch-proteome profiling, and following the steps below will enable users to properly execute this approach.

**CRITICAL:** There are numerous ways these steps can be fine-tuned to fit the equipment available to the lab and the sample types to be assessed in a given experiment or scientific question. We encourage each project team to optimize this approach based on their own equipment and experimental set ups.

1. Digestion into Tryptic Peptides for LC-MS

a. Dispense 200nL of 40ug/mL trypsin in 60mM TEAB (triethylammonium bicarbonate) into each plate well to digest the cell lysates.
b. Incubate the 384-well plate at 37 C for 4 hrs
c. c. Quench the plate with 200nL 0.1% formic acid solution.
d. Spin plate at 500 rcf Pause Point: Cover plate and freeze at -80C until ready for LC MS acquisition
e. Place the plate on an autosampler for LC MS
2. LC-MS Data Acquisition Using nanoDTSC
  a. QC Monitoring of LC-MS System: Ensure instrument performance by analyzing 1 ng of HeLa in at least duplicate using the standard method. For this protocol approximately 4,000 protein identifications indicate adequate instrument performance. Space blank runs at the beginning, end and regular intervals (e.g., every 20-50 cells) throughout acquisition run to ensure low carryover.
  Note: In the method described, we perform LC-MS analysis on an Ultimate nanoRSLC-3500 (Dionex/Thermo) liquid chromatography systems (LC), configured for nanoscale dual-trap single column reversed phase chromatography and a timsTOF Ultra 2 mass spectrometer (nanoDTSC; see Kreimer, 2023).
  a. Configure the LC with one 6-port and one 10-port valve within the oven compartment
  b. Separate high flow Ultimate 3000 pump for cleaning and loading the trapping columns.
  c. Install in the 10-port valve two Exp 2 stem trapping columns (Optimize Technologies) packed with 170 nL of 10 µm PLRP solid phase (Agilent)
  d. While the analytical gradient flowed through one trapping column into the analytical column (PepSep 15 cm x 75 um packed with 1.9 um C18 Reprosil AQ stationary phase from Bruker) and electrosprayed through a 10 um CaptiveSpray emitter (Bruker), the alternate trapping column would be washed with organic (acetonitrile, 0.1% formic acid) solvent equilibrated with water 0.1% formic acid and loaded with the next sample
  e. Switch the position of the 10-port valve to start analysis of the loaded sample with no delay
  f. The 6-port valve dictates the washing and loading direction Note: This parallelization enables sample resuspension with the autosampler without effect instrument down time, a limitation of conventional trap and elute methods
  g. Ionize chromatographic eluate and analyze by Mass Spectrometry (MS). In this example, the Ultimate3000 LC-MS nano-DualTrap system is interfaced to a Bruker TIMSTOF Ultra2 Mass Spectrometer operating under the following specifications:
    i. Achieve positive electrospray ionization with 1600 V potential on the capillary and dry nitrogen gas at 200 C and 3.0 L/min flowrate.
    ii. Acquire data using a Data Independent Acquisition Parallel Accumulation Serial Fragmentation (dia-PASEF) method with a 0.96 s cycle time and 100 ms ramp and accumulation time, and the MS1 isolation windows
    Note: Acquire three or more runs of 5-10ng additional digest of bulk lysate representing the cell types / tissue of interest on the same set up, using matching LC and either DIA or Data Dependent Acquisition (DDA) mode (or both) to be used as a sample specific peptide library for downstream bioinformatic peptide sequence assignment.
3. Peptide Sequence Identification and Protein Quantification Estimation Note: Many options are available for bioinformatics processing of LC-MS DIA datasets. For the following steps, we leverage a sample specific peptide library which we generate from bulk mouse brain lysates from nine UM-HET3 mice (Miller et al., 1997; Moore & Murphy, 2026; Warner et al., 2000) (See Expected Outcome; Figure 5). We digest lysates with trypsin and analyze using matching LC-MS set up relative to our individual cell proteomes. Library free searching against a species-specific sequence database, or use of generic, sample-representative library using publicly available datasets generated on a similar instrument used for the experiment are options for peptide sequence assignment. In our hands, we often find these two latter options generate lower peptide and protein group matches per cell relative to the sample-specific library approach. Users should explore and optimize ideal methods for their respective biological samples. The following describes the method we apply in this study:
  a. Search raw files in DIA Neural Networks (DIA-NN) software (in this case version 1.8.1, but the most recent stable release is appropriate for new studies; Demichev et al., 2020).
  b. Use a sample specific peptide library. Libraries made using DIA can be generated directly using a library free search in DIA-NN according to published instructions. Libraries made using DDA runs can be assembled using the FragPipe workflow, again following published instructions. Load the resulting library file into DIA-NN for single-cell analysis.
  c. Set match between runs (MBR) to ‘on’ and reannotation of gene groups to a supplied mouse FASTA protein sequence database
  d. Do not apply heuristic protein inference and turn off all intensity normalization (this will be achieved separately in later data processing steps)
  e. Produce protein quantification estimates using the DIA-NN maxLFQ algorithm Note: DIA-NN outputs several processed protein quantification matrices, with sample identities in columns and the protein group (.pg file), gene group (.gg file), or peptide (.pr file) in rows (See MassIVE raw dataset).
  f. Use the protein group output matrix and remove all instances of proteins identified by shared peptide sequences across more than one gene (referred to as “non-proteotypic”, see Exclusion Criteria; Table 1). This protein-by-cell matrix now only includes proteins quantified by proteotypic peptides in the final dataset.
4. Pre-processing of Single Cell Proteomic Data These steps will refine the protein-by-cell matrix from DIA-NN for downstream analysis (e.g. correlation to other cellular measures like electrophysiology or morphology, differential expression analysis, etc.).
  a. For pre-processing (data imputation, exclusion, normalization, transformation, and annotation), use the Scanpy python (version 1.11.5, or other similar R or python) package, which is publicly available through the Python Package Index (PyPI) and can be installed using pip (the Python package installer).
    i. Upload the protein-by-cell data matrix from DIA-NN in CSV format and convert it into an annotated data matrix (AnnData) object using the Scanpy single cell data analysis workflow.
    ii. Impute missing protein abundance values as zero (under the assumption that non-detected proteins represented absent expression).
    iii. Exclude proteins detected in fewer samples than required by a user-defined cutoff criterion. Note: The precise cutoff criterion will depend on the goals of the particular study or scientific question. Statistical analysis requires detection in at least 3 samples, so n=3 can be used as the most relaxed threshold for exclusion, which is appropriate for discovery or exploratory experiments. For hypothesis-testing and mechanistic studies, a more stringent criterion may be recommended (e.g. protein is detected in at least 50% of the samples).
    iv. To make data comparable across samples, normalize individual protein intensities to the total protein abundance for each sample.
    v. To reduce the effect of very high abundance proteins on the overall analysis, scale each normalized protein abundance using a natural logarithm transformation. Note: as an optional step, update imputed values for undetected proteins (entered as a “0” value) using linear regression to predict non-zero values. This step can be used to predict values for undetected proteins based on measured values from other samples in the data set, thereby functionally increasing sample size for increased statistical power in downstream analyses.
  b. Perform dimensionality reduction, data visualization, and batch correction (if necessary).
    i. Use the Principle Component Analysis (PCA) function in Scanpy to reduce dimensionality of proteomic data.
    ii. Perform clustering analysis (e.g. Leiden-based clustering in Scanpy) to analyze similarity between each sample and determine level of batch effect.
      1. Use default parameters except set resolution to 1.2
      2. The Davies-Bouldin Index (DBI) metric reflects the level of separation/clustering of samples. Typical ranges are defined as: <0.5 = excellent separation, high clustering; 0.5 – 1 = moderate clustering; 1 – 2 = moderate clustering; >2 = substantial overlap and low clustering.
    iii. Use Uniform Manifold Approximation and Projection (UMAP) in Scanpy for data visualization and exploration. Note: If the data strongly cluster based on non-biologically relevant variables (e.g. from which plate the sample was measured), it is recommended to perform batch effect correction using relevant packages/software (e.g. ComBat-Seq in R). Note: Optionally, add cell type annotation using canonical marker genes (e.g. MapMyCells from Allen Brain Institute).

**Table 1.** Protein assignment based on proteotypic and non-proteotypic peptides.

| Identified Peptide | Corresponding Protein | Classification | Application |
| --- | --- | --- | --- |
| Peptide A | Protein X | Proteotypic | Correlative Analysis |
| Peptide A<br>Peptide B<br>Peptide C | Protein X | Proteotypic | Correlative Analysis |
| Peptide A | Protein X; Protein Y | Non-proteotypic | Descriptive analysis |

After completion of the above steps, you will have a protein-by-cell matrix which is appropriate for downstream analysis, such as association to additional cellular parameters (e.g. electrophysiological or morphological), differential expression, or descriptive proteomics (see Fig 4 for overall workflow).

**Figure 4.**
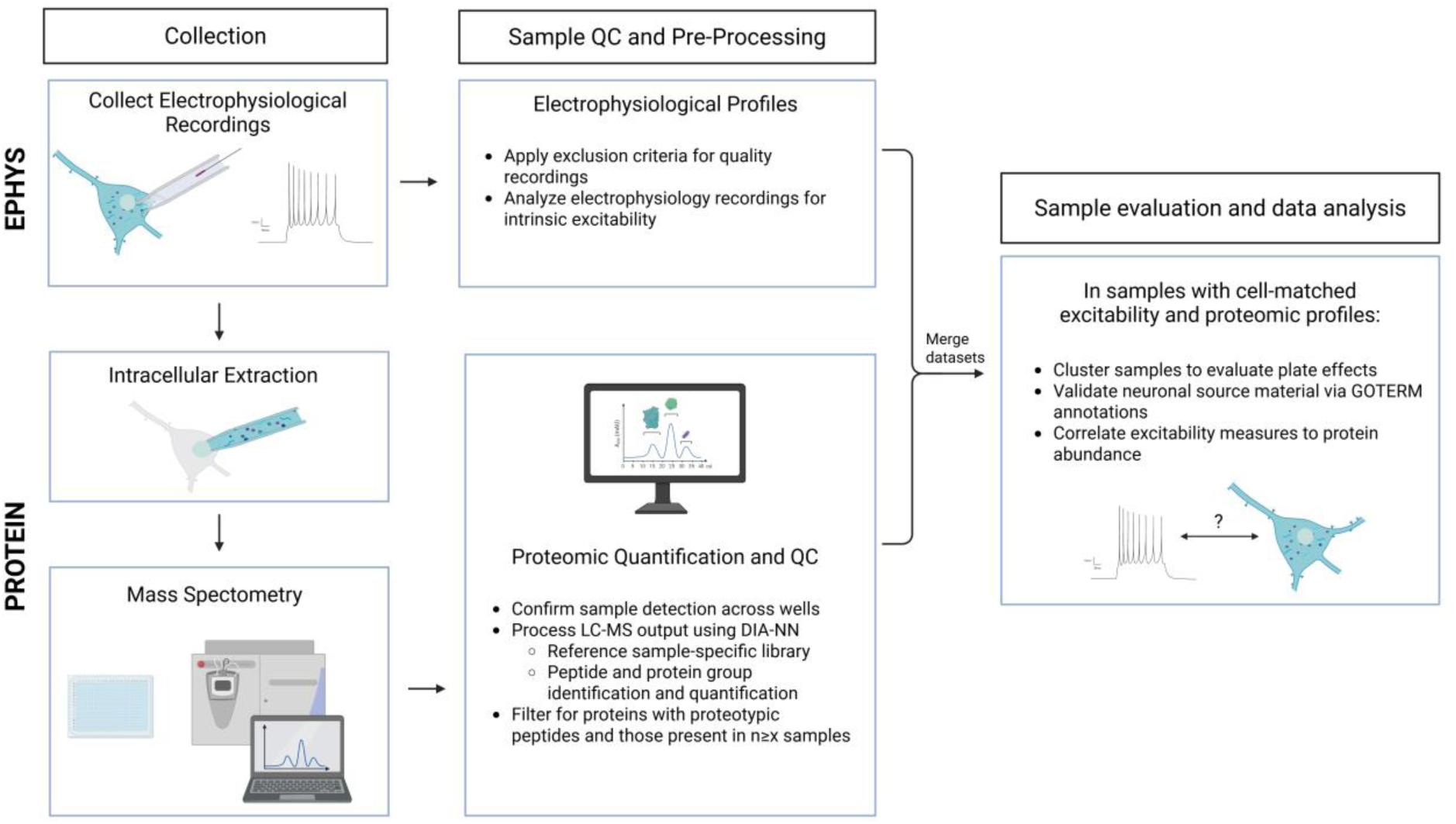
Patch-proteomics sample collection and data processing workflow. Electrophysiological recordings are collected during whole-cell patch-clamp, followed by the extraction of intracellular contents. This extracted sample is expelled into a 384-well plate and saved for LC MS. For electrophysiological recordings, apply exclusion criteria to ensure high quality recordings and measure properties of intrinsic excitability or other neurophysiological features (EPHYS, top). For extracted samples, confirm peptide detection across all wells and process the raw MS data using DIA-NN software. Reference raw output against a sample-specific library for peptide and protein identification and quantification. Filtering for proteins with proteotypic peptides and proteins present in a minimum number of samples will further ensure reliable interpretation of proteomic profiles (PROTEIN, bottom). After QC, merge electrophysiological and proteomic measurements into a unified, cell-matched dataset. Evaluate the effects of multiple plates through cluster analysis to ensure technical consistency across all plates. Functional annotation of protein list via DAVID analysis validates source material and provides additional characterization of the collected proteomes. Associate electrophysiological measures and protein abundance by performing linear regression analysis (Sample evaluation and data analysis, right).

### Expected outcomes

The results described in this section are from a patch-proteomics benchmark study to develop and validate this pipeline. These data demonstrate the capacity to reliably identify and quantify peptides and their corresponding proteins from individual neurons. We highlight the results by utilizing a sample-specific library from genetically diverse UM-HET3 mice that simulate population variability across mouse strains (Miller et al., 1997; Moore & Murphy, 2026; Warner et al., 2000). When following the above protocol, this technique can be successfully implemented to characterize proteomes within different brain regions, cell types, and physiological, pathological, and behavioral contexts.

#### Benchmark single-cell patch proteomics study

We perform whole-cell patch-clamp on pyramidal neurons in acute hippocampal slices from mice, generating electrophysiological profiles from 155 subicular neurons in 44 UM-HET3 mice, comprising adult (11-13 month) and aged (26-28 month) females and males (Figure 5, top left). At the conclusion of the recording session for each neuron, we extract and collect intracellular samples, which we successfully complete in 117 out of 155 patched neurons (Figure 5, bottom left). For each electrophysiological recording, we apply QC parameters which removes low-quality recordings, interneurons, and burst-spiking (BS) cells. This leaves 100 regular-spiking (RS) pyramidal neurons for analysis of intrinsic excitability properties, including Resting Membrane Potential (RMP), Input Resistance (Rin), sag ratio, firing frequency, properties of single action potentials (AP), Rheobase, and the medium and slow post-burst afterhyperpolarization (mAHP, sAHP) (Figure 5, top-middle; Supplemental Table S1). From the 117 intracellular samples we successfully collect, LC-MS detects peptides in 102 samples. Across this entire population, we identify and quantify a total of 13,285 peptides (4,435 ± 248 peptides per sample), 2,393 protein groups, and 2,359 proteotypic proteins (1,142 ± 48.1 proteotypic proteins per sample). We further exclude any peptides/proteins that we detect in fewer than three samples, which results in a total of 2,237 proteins for proteomic characterization (Figure 5A-C; see Exclusion Criteria).

**Figure 5.**
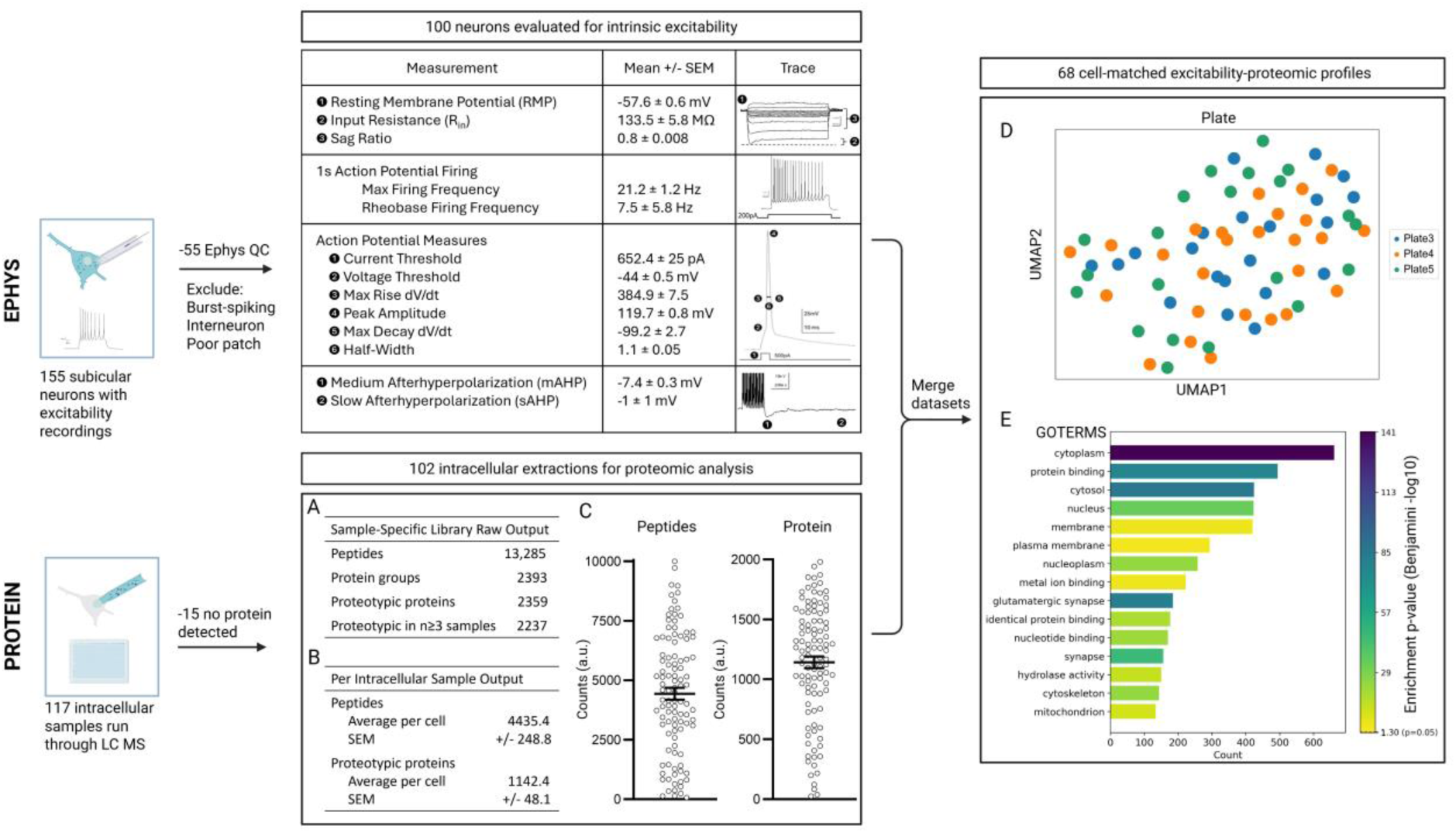
Patch-proteomics benchmark study: data processing and output. In this study, 155 electrophysiological profiles and 117 intracellular samples were collected (EPHYS, PROTEIN; left). For electrophysiology characterization, we exclude 55 samples for quality control purposes and cell type (see Exclusion Criteria) leaving 100 electrophysiological samples, each containing a series of recordings we use to evaluate intrinsic excitability (see table, top-middle). For proteomic characterization, we exclude 15 of the 117 samples due to no peptide detection or failed injection during LC MS. A) When employing a sample specific peptide library, the remaining 102 samples result in an output of 13,285 identified peptides, 2,393 proteins groups, 2,359 proteotypic proteins, and 2,237 proteotypic in at least three samples. B-C) On average, there are 4,435 ± 248 peptides per sample and 1,142 ± 48.1 proteins (from proteotypic peptides) per sample. D) Cluster analysis of proteomic profiles shows no plate effect across the three well plates used for this study. E) GOTERM annotation using DAVID shows strong cytoplasmic, nuclear, and neuron specific proteins.

Following this data QC, processing, and analysis for both electrophysiological and proteomic measurements, we merge these data sets into 68 cell-matched profiles for downstream correlational analysis (Supplemental Table S2). We first ensure that proteomic samples collected over time and measured from different plates represent an assessment of the same independent population of proteins by visualizing clustering based on technical (i.e. sample plate), rather than biological factors. Our uniform manifold approximation and projection (UMAP) visualization shows that proteomic samples from different plates have low clustering/high overlap and thus represent a clear example of successful technical integration of samples across multiple plates (Figure 5D).

We next examine these proteins in terms of their cellular compartment and functional annotations to evaluate and validate the specific proteomic profiles we detect in our neuronal population. Utilizing the Database for Annotation, Visualization, and Integrated Discovery (DAVID; Huang et al., 2009; Sherman et al., 2022), we functionally annotate the list of proteins using GOTERMs (GOTERM_DIRECT_MF, GOTERM_DIRECT_BP, GOTERM_DIRECT_CC). As expected, this analysis shows that there is a robust enrichment of cytoplasmic and nuclear proteins from our samples, along with mitochondrial, cytoskeleton, and plasma membrane proteins. Importantly, synaptic and neuron-specific proteins are prominently represented in these functional annotations, confirming our specific interrogation of the neuronal proteome (top 15 categories represented shown in Figure 5E; for complete DAVID chart of all GOTERMS, see Supplemental Table S3).

To uncover relationships between proteins and neuronal function, we perform a linear regression (see Quantification and Statistical Analyses) that identifies associations between electrophysiological properties and protein abundance, with each electrophysiological measurement modeled as the outcome variable, protein abundance as the predictor, and sex and age included as covariates (Supplemental Table S4). We first probe these correlational data to validate this paradigm by identifying known protein-electrophysiology relationships which have been implicated in prior work.

For instance, previous studies have shown that increased excitability in hippocampal neurons is associated with learning and memory, even in aging and AD mouse models (Kaczorowski et al., 2011; Kaczorowski & Disterhoft, 2009). As expected, our analysis identifies proteins known to affect neuronal excitability, including ATPase-dependent pumps, ion channels, and glutamate receptor complexes as being associated with multiple measures of intrinsic excitability in our electrophysiological recordings (Supplemental Table S6). Notably, the Na+/K+-ATPase pump subunit AT1A3, which has a vital role in maintaining electrochemical gradients, correlates with multiple measures of excitability (negatively correlated with R_in_; positively correlated with Rheobase and I_threshold_), and indicates that greater expression of this protein is associated with lower excitability. Interestingly, and consistent with our results, recent work shows that dysfunction of this specific subunit leads to hyperexcitability, which further substantiates and validates our patch-proteomics approach to detect functional proteomic changes that mediate neuronal extricability (Akkuratov et al., 2025; Skou & Esmann, 1992).

In addition to expected proteins, we also discover novel proteins that strongly correlate with electrophysiological properties and thus represent intriguing, nominated candidates to investigate for their roles in neuronal excitability (Supplemental Table S6). These results highlight the incredible utility and power that patch-proteomics provides for discovery biology. Thus, leveraging this paradigm will enable an unbiased, single-neuron approach to reveal novel proteomic signatures relevant to learning and memory, normal aging, and neurodegenerative disease.

### Quantification and statistical analysis

#### Exclusion Criteria

Filtering and QC parameters for both electrophysiological and proteome data ultimately depend on the specific questions of interest in the experiment being conducted. For example, electrophysiological data needs to be filtered based on cell type (e.g. pyramidal neuron vs interneuron) as well as assessed for “high-quality” recording.

In the data presented here, we choose to investigate subicular pyramidal regular-spiking neurons. These are not thought to be spontaneously active, so as a measure of cell and recording quality we employ a physiologically relevant exclusion criterion in which the resting membrane potential (RMP) is required to be below (more negative) than the voltage threshold of a given cell to be acceptable for analysis. To exclude burst-spiking (BS) cells, we measure single APs following a 2 ms current injection. Cells that have two (or more) APs riding on the same depolarizing envelope with an instantaneous frequency of >100 Hz are classified as BS (Dunn et al., 2018). To exclude interneurons, we measure firing frequency in response to a 1-s step current injection. Cells that maintain a very high frequency (>100 Hz) of firing throughout the entire step (i.e. do not accommodate) and have half-widths <1 ms are classified as interneurons.

An important QC step for proteomic data is to define exclusion criteria based on peptide-to-protein correspondence. A single peptide fragment identified by LC-MS can either map uniquely to a single protein (which we refer to as “proteotypic”) or can map to multiple proteins, which cannot be uniquely resolved (referred to as “non-proteotypic”). Additionally, multiple peptides can uniquely correspond to a single protein, which is also considered proteotypic. In the case of non-proteotypic peptides, the specific protein detected by LC-MS cannot be resolved from the sequence, so we exclude these proteins from downstream analysis to focus on specific, identified protein-excitability relationships. It is important to note, however, that these non-proteotypic peptides often correspond to similar types of proteins (e.g. Different, yet closely related histone subunits), and even though a specific protein cannot be deciphered, users may decide to include these non-proteotypic groups as descriptive data in their characterization of their proteomic datasets (Table 1). In addition, it is important to consider whether a cut-off for the frequency of protein observation will be used (i.e. in how many samples a given protein is detected). In the data presented here, we maximize proteomic coverage for discovery-based analysis by using the most liberal cutoff that still retains the ability for statistical analysis, which is that a protein need only be detected in a minimum of three independent samples. Investigators may choose a more stringent exclusion criterion (e.g. protein present in >50% of samples) if focusing on more ubiquitously expressed proteins.

#### Linear Regression Analysis of Protein–Electrophysiology Associations

To evaluate associations between protein abundance and electrophysiological phenotypes while controlling potential confounding variables, we perform linear regression separately for each protein. To account for the potential effects of the sex and age of the animals from which our samples, we use a reduced linear model, containing only covariates (ephys ∼ sex + age), to determine the effect of these variables alone on electrophysiological outcomes (quantified by the “goodness of fit” adjusted R-squared value [R^2^_adj_]). We then compare this result to that obtained using a full linear model which includes the protein term (ephys ∼ protein + sex + age). The magnitude of the effect of each protein (i.e. increase in variance explained beyond covariate alone) is then quantified as the change in the R^2^_adj_ (ΔR^2^_adj_), calculated by subtracting the R^2^_adj_ of the reduced model from the R^2^_adj_ of the full model. To incorporate the directionality of the effect of each given protein on electrophysiological outcomes, we multiply the (non-negative) ΔR^2^_adj_ values by the sign of the protein regression coefficient (β) from the full linear model; for simplicity, this signed ΔR^2^_adj_ metric is referred to as the overall “Adjusted R^2^” (note capital [“Adjusted”] to distinguish from lowercase “adjusted” referring to metric from individual linear models). Finally, we calculate statistical significance for each protein-by-electrophysiological outcome using the Benjamini–Hochberg multiple testing correction false discovery rate procedure

To visualize the effect of individual proteins on electrophysiological outcomes, we generate a volcano plot where Adjusted R^2^ is plotted against the -log10(p) for each protein-electrophysiological measure pair (Figure 6). Thus, positive associations where larger values of the electrophysiological measurement correspond to increased protein expression are displayed on the positive x-axis, while those where larger electrophysiological values correspond to decreased protein expression are displayed on the negative x-axis.

**Figure 6.**
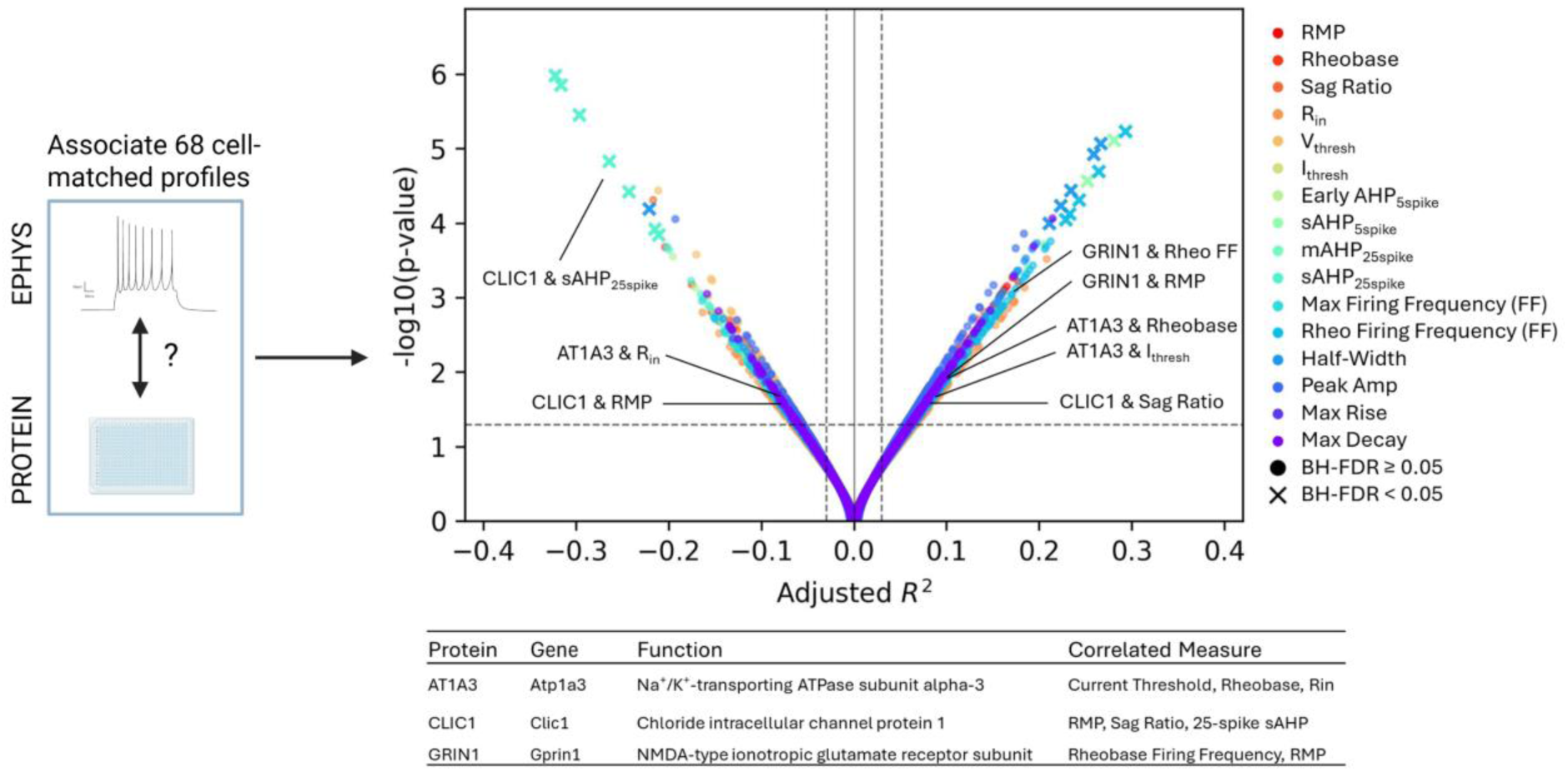
Cell-matched profiles reveal specific proteins associated with properties of intrinsic excitability. Following quality control and data processing, linear regression is performed to associate protein abundance with each measure of intrinsic excitability. Numerous proteins with known roles in excitability and plasticity significantly associate with multiple excitability properties (AT1A3, CLIC1, GRIN1). Many novel proteins strongly associate with electrophysiological properties and represent interesting candidates to further investigate roles in neuronal excitability.

Interestingly, and in support of the validity and power of this approach, we find expected associations between electrophysiological outcomes and ionic channels, receptors, and transporters, including CLIC1 (a chloride intracellular channel protein), GRIN1 (a receptor subunit of the NMDA-type ionotropic glutamatergic receptor), and AT1A3 (a ATP-dependent subunit of the Na+/K+-transporter). Taken together, this analysis reveals 1,078 proteins significantly associated with electrophysiological outcomes (Figure 6). Of these, 20 proteins remain significant after Benjamini-Hochberg (BH) correction (denoted with an “X” in Figure 6), which represent potential novel targets to mediate neuronal excitability changes associated with learning and memory, aging, and/or disease states.

## Limitations

Intracellular sample collection is done by suctioning the cytosol and the nucleus from the soma into the recording pipette. This means proteins that exist in the distal axons and dendrites will not be collected. However, this does not mean that synaptic proteins will not be detected. Many critical synaptic proteins are translated in the soma and trafficked to the distal processes (Okada et al., 2009; Sorokina et al., 2026). This is evident from the patch-proteomics study highlighted in this protocol, as one of the top functional annotations for detected proteins was “synapse”. This indicates that despite missing out on synaptically localized proteins, the capacity to detect newly synthesized or somatically localized versions of these proteins is apparent.

## Troubleshooting

### Problem 1

Loss of whole-cell patch during extraction

### Potential solution

Apply less negative pressure. The amount required will vary from cell to cell, and depends on several factors:

a. Quality of patch
b. Cell type
c. Cell health
d. Size of cell
e. Pipette resistance
f. Specifications of the tubing, syringes, connectors, and valves of the patching setup Note: Try a variety of strengths of negative pressure to see what works best. Every cell and every patch is a little different. Repetition and trial and error are key to gaining experience to know what pressure is right. Note: the negative pressure applied during extraction may lower the access resistance (R_a_) due to clearing any cellular debris from the pipette opening. But it should not be enough to destabilize and lose the whole-cell patch.

### Problem 2

Unable to retract nucleus from cell

### Potential solutions

1. Ensure the pipette does not drift away from the original patch position on the soma throughout the recordings.
2. Apply stronger negative pressure during extraction phase
3. Spend more time locating and extracting nucleus
4. Slow down the pace of retraction to allow for cell membrane to relax and “release” the nucleus (Figure 7.)

### Problem 3

Unable to achieve a 1 GΩ seal on nucleus upon retraction

**Figure 7.**
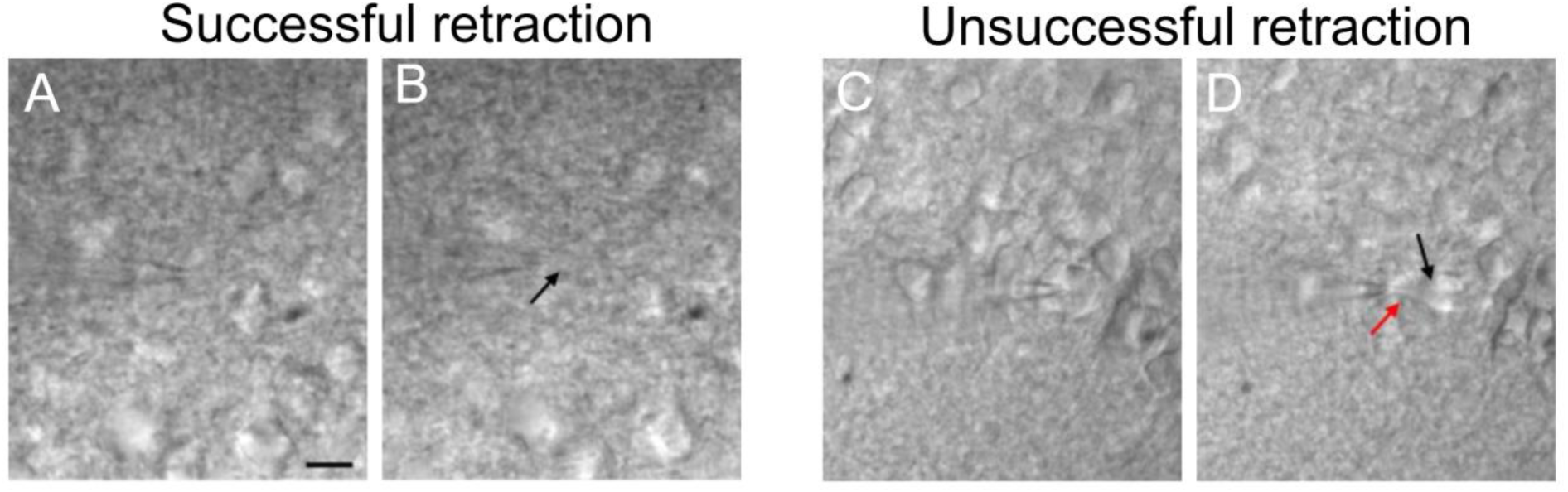
Successful versus unsuccessful retraction of the cytosol and nucleus. A) Patched neuron prior to successful retraction. B) Successful retraction with the oval-shaped nucleus following the pipette (black arrow = nucleus). C) Patched neuron prior to unsuccessful retraction of the nucleus. D) Unsuccessful retraction, with the cell membrane stretching (red arrow) and nucleus being left behind in the soma (black arrow; compare to B). (Scale bar = 10uM)

### Potential solutions

1. Spend more time in the extraction phase attempting to attach the nucleus to the end of the pipette
2. Slow down the pace of retraction to allow for cell membrane to “release” the nucleus Note: Regardless of the R_m_ achieved during retraction, if the nucleus is visually attached to the end of the pipette tip, we recommend saving the sample and keeping detailed notes of whether a GΩ nucleated patch occurred.

### Problem 4

Low protein yield per cell

### Potential solution

1. One cause may be digestion efficiency. This can be assessed through quantifying the percentage of missed trypic cleavages among the peptides that were identified or through inclusion of a spiked digestion standard protein like e.coli beta galactosidase to monitor trypsin performance. If digestion is sub-optimal:

a. Check digestion buffer pH and ensure a pH > 7, ideally between 7-9
b. Change digestion buffer to 10 mM CaCl2 in 50mM TRIS
2. Add more Trypsin or combine Trypsin and Lys-C enzymes
  a. Another cause could be LC-MS performance
3. Check protein IDs from HeLa QC runs and MS calibration, if low assess the trap and column performance and replace as necessary, or consider cleaning ion transfer tube or other MS components
  a. Avoid utilizing library free or generic libraries for bioinformatics analysis and generate a sample specific library per this protocol recommendation

## Supporting information

Sup Table 1 - Excitability Measures

Sup Table 2 - Excitability and Protein by Sample

Sup Table 3 - DAVID annotations

Sup Table 4 - Excitability to Protein associations

## Resource availability

### Lead contact

Further information and requests for resources and reagents should be directed to and will be fulfilled by the lead contact, Dr. Catherine C. Kaczorowski

### Technical contact

Electrophysiology and intracellular extraction: J. Christian Althaus

LC-MS and bioinformatics: Sarah J. Parker

Downstream data analysis: Yiding Cao

### Materials availability

This study did not generate any new unique reagents and/or materials

### Data and code availability

The mass spectrometry proteomics data have been deposited to the MassIVE Data Repository (https://massive.ucsd.edu/) under the dataset identifier MSV000102490.

## Acknowledgments

This work was supported by the National Institute on Aging R01 AG075818 (C.C.K.) and the Simons Foundation, Simons Collaboration on Plasticity and the Aging Brain Research Award 984481 (C.C.K.).

We would like to thank and acknowledge the Cedars Sinai Proteomics and Metabolomics Core facility for their effort in generating these data.

## Author contributions

Conceptualization, C.C.K.; Methodology, J.C.A., Y.C., S.J.M., S.J.P., and C.C.K; Investigation, J.C.A., K.S., A.H., and R.C.; Writing – Original Draft, J.C.A., Y.C., and S.J.P; Writing – Review & Editing, S.J.M., and C.C.K. ; Supervision, S.J.M, S.J.P., and C.C.K. Funding Acquisition, S.J.P. and C.C.K.; Resources, Y.C., K.S., A.H., and R.C.

## References

Bubis, J. A., Arrey, T. N., Damoc, E., Delanghe, B., Slovakova, J., Sommer, T. M., Kagawa, H., Pichler, P., Rivron, N., Mechtler, K., & Matzinger, M. (2025). Challenging the Astral mass analyzer to quantify up to 5,300 proteins per single cell at unseen accuracy to uncover cellular heterogeneity. Nature Methods, 22(3), 510–519. 10.1038/s41592-024-02559-1

Cadwell, C. R., Palasantza, A., Jiang, X., Berens, P., Deng, Q., Yilmaz, M., Reimer, J., Shen, S., Bethge, M., Tolias, K. F., Sandberg, R., & Tolias, A. S. (2016). Electrophysiological, transcriptomic and morphologic profiling of single neurons using Patch-seq. Nature Biotechnology, 34(2), 199–203. 10.1038/nbt.3445

Cadwell, C. R., Scala, F., Li, S., Livrizzi, G., Shen, S., Sandberg, R., Jiang, X., & Tolias, A. S. (2017). Multimodal profiling of single-cell morphology, electrophysiology, and gene expression using Patch-seq. Nature Protocols, 12(12), 2531–2553. 10.1038/nprot.2017.120

Chen, H., Shi, X., & Guo, F. (2025). Whole-cell patch clamp and extracellular electrophysiology recordings in mouse brain slices. STAR Protocols, 6(3). 10.1016/j.xpro.2025.104008

Ctortecka, C., Clark, N. M., Boyle, B., Seth, A., Mani, D. R., Udeshi, N. D., & Carr, S. A. (2024). Automated single-cell proteomics providing sufficient proteome depth to study complex biology beyond cell type classifications. bioRxiv: The Preprint Server for Biology, 2024.01.20.576369. 10.1101/2024.01.20.576369

Demichev, V., Messner, C. B., Vernardis, S. I., Lilley, K. S., & Ralser, M. (2020). DIA-NN: Neural networks and interference correction enable deep proteome coverage in high throughput. Nature Methods, 17(1), 41–44. 10.1038/s41592-019-0638-x

Deribe, Y. L., Pawson, T., & Dikic, I. (2010). Post-translational modifications in signal integration. Nature Structural & Molecular Biology, 17(6), 666–672. 10.1038/nsmb.1842

Dunn, A. R., Neuner, S. M., Ding, S., Hope, K. A., O’Connell, K. M. S., & Kaczorowski, C. C. (2018). Cell-type-specific changes in intrinsic excitability in the subiculum following learning and exposure to novel environmental contexts. eNeuro, 5(6). 10.1523/ENEURO.0484-18.2018

Graves, A. R., Moore, S. J., Spruston, N., Tryba, A. K., & Kaczorowski, C. C. (2016). Brain-derived neurotrophic factor differentially modulates excitability of two classes of hippocampal output neurons. Journal of Neurophysiology, 116(2), 466–471. 10.1152/jn.00186.2016

Hartl, F. U., Bracher, A., & Hayer-Hartl, M. (2011). Molecular chaperones in protein folding and proteostasis. Nature, 475(7356), 324–332. 10.1038/nature10317

Huang, D. W., Sherman, B. T., & Lempicki, R. A. (2009). Systematic and integrative analysis of large gene lists using DAVID bioinformatics resources. Nature Protocols, 4(1), 44–57. 10.1038/nprot.2008.211

Jarsky, T., Mady, R., Kennedy, B., & Spruston, N. (2008). Distribution of bursting neurons in the CA1 region and the subiculum of the rat hippocampus. The Journal of Comparative Neurology, 506(4), 535–547. 10.1002/cne.21564

Josselyn, S. A., & Tonegawa, S. (2020). Memory engrams: Recalling the past and imagining the future. Science, 367(6473), eaaw4325. 10.1126/science.aaw4325

Kaczorowski, C. C., Davis, S. J., & Moyer, J. R. (2012). Aging redistributes medial prefrontal neuronal excitability and impedes extinction of trace fear conditioning. Neurobiology of Aging, 33(8), 1744–1757. 10.1016/j.neurobiolaging.2011.03.020

Kaczorowski, C. C., & Disterhoft, J. F. (2009). Memory deficits are associated with impaired ability to modulate neuronal excitability in middle-aged mice. Learning & Memory, 16(6), 362–366. 10.1101/lm.1365609

Kaczorowski, C. C., Sametsky, E., Shah, S., Vassar, R., & Disterhoft, J. F. (2011). Mechanisms underlying basal and learning-related intrinsic excitability in a mouse model of Alzheimer’s disease. Neurobiology of Aging, 32(8), 1452–1465. 10.1016/j.neurobiolaging.2009.09.003

Kreimer, S., Binek, A., Chazarin, B., Cho, J. H., Haghani, A., Hutton, A., Marbán, E., Mastali, M., Meyer, J. G., Mesquita, T., Song, Y., Van Eyk, J., & Parker, S. (2023). High-Throughput Single-Cell Proteomic Analysis of Organ-Derived Heterogeneous Cell Populations by Nanoflow Dual-Trap Single-Column Liquid Chromatography. Analytical Chemistry, 95(24), 9145–9150. 10.1021/acs.analchem.3c00213

Lee, B. R., Budzillo, A., Hadley, K., Miller, J. A., Jarsky, T., Baker, K., Hill, D., Kim, L., Mann, R., Ng, L., Oldre, A., Rajanbabu, R., Trinh, J., Vargas, S., Braun, T., Dalley, R. A., Gouwens, N. W., Kalmbach, B. E., Kim, T. K., … Berg, J. (2021). Scaled, high fidelity electrophysiological, morphological, and transcriptomic cell characterization. eLife, 10, e65482. 10.7554/eLife.65482

Lipovsek, M., Browne, L., & Grubb, M. S. (2020). Protocol for Patch-Seq of Small Interneurons. STAR Protocols, 1(3), 100146. 10.1016/j.xpro.2020.100146

Liu, Y., Beyer, A., & Aebersold, R. (2016). On the Dependency of Cellular Protein Levels on mRNA Abundance. Cell, 165(3), 535–550. 10.1016/j.cell.2016.03.014

Miller, R. A., Chrisp, C., & Galecki, A. (1997). CD4 memory T cell levels predict life span in genetically heterogeneous mice. The FASEB Journal, 11(10), 775–783. 10.1096/fasebj.11.10.9271362

Moore, S. J., & Murphy, G. G. (2026). Acarbose improves cognitive function in a mouse model of normal aging but not Alzheimer’s disease (p. 2026.04.28.721469). bioRxiv. 10.64898/2026.04.28.721469

Nakai, K., Horton, P., Nakai, K., & Horton, P. (1999). PSORT: A program for detecting sorting signals in proteins and predicting their subcellular localization. Trends in Biochemical Sciences, 24(1), 34–35. 10.1016/S0968-0004(98)01336-X

Okada, D., Ozawa, F., & Inokuchi, K. (2009). Input-Specific Spine Entry of Soma-Derived Vesl-1S Protein Conforms to Synaptic Tagging. Science, 324(5929), 904–909. 10.1126/science.1171498

Pascal, L. E., True, L. D., Campbell, D. S., Deutsch, E. W., Risk, M., Coleman, I. M., Eichner, L. J., Nelson, P. S., & Liu, A. Y. (2008). Correlation of mRNA and protein levels: Cell type-specific gene expression of cluster designation antigens in the prostate. BMC Genomics, 9, 246. 10.1186/1471-2164-9-246

Saddic, L., Kaneda, G., Momenzadeh, A., Zilberberg, L., Song, Y., Mastali, M., Kreimer, S., Hutton, A., Haghani, A., Meyer, J., & Parker, S. (2025). Single Cell Proteomics Reveals Novel Cell Phenotypes in Marfan Mouse Aneurysm. Systems Biology. 10.1101/2025.02.15.638465

Schimanski, L. A., & Barnes, C. A. (2010). Neural Protein Synthesis during Aging: Effects on Plasticity and Memory. Frontiers in Aging Neuroscience, 2. 10.3389/fnagi.2010.00026

Schwanhäusser, B., Busse, D., Li, N., Dittmar, G., Schuchhardt, J., Wolf, J., Chen, W., & Selbach, M. (2011). Global quantification of mammalian gene expression control. Nature, 473(7347), 337–342. 10.1038/nature10098

Sherman, B. T., Hao, M., Qiu, J., Jiao, X., Baseler, M. W., Lane, H. C., Imamichi, T., & Chang, W. (2022). DAVID: A web server for functional enrichment analysis and functional annotation of gene lists (2021 update). Nucleic Acids Research, 50(W1), W216– W221. 10.1093/nar/gkac194

Sorokina, O., Bulovaite, E., Sorokin, A., Grant, S. G. N., & Armstrong, J. D. (2026). Protein trafficking and synaptic demand configure complex and dynamic synaptome architectures of individual neurons. Scientific Reports, 16(1), 11541. 10.1038/s41598-026-40513-7

Tasaki, S., Xu, J., Avey, D. R., Johnson, L., Petyuk, V. A., Dawe, R. J., Bennett, D. A., Wang, Y., & Gaiteri, C. (2022). Inferring protein expression changes from mRNA in Alzheimer’s dementia using deep neural networks. Nature Communications, 13(1), 655. 10.1038/s41467-022-28280-1

Upadhya, S. R., & Ryan, C. J. (2022). Experimental reproducibility limits the correlation between mRNA and protein abundances in tumor proteomic profiles. Cell Reports Methods, 2(9), 100288. 10.1016/j.crmeth.2022.100288

Vogel, C., & Marcotte, E. M. (2012). Insights into the regulation of protein abundance from proteomic and transcriptomic analyses. Nature Reviews Genetics, 13(4), 227–232. 10.1038/nrg3185

Walker, H., & Frost, N. A. (2024). Distinct transcriptional programs define a heterogeneous neuronal ensemble for social interaction. iScience, 27(7), 110355. 10.1016/j.isci.2024.110355

Wang, X., Pattison, J. S., & Su, H. (2013). Post-translational Modification and Quality Control. Circulation Research, 112(2), 367–381. 10.1161/CIRCRESAHA.112.268706

Warner, H. R., Ingram, D., Miller, R. A., Nadon, N. L., & Richardson, A. G. (2000). Program for testing biological interventions to promote healthy aging. Mechanisms of Ageing and Development, 115(3), 199–207. 10.1016/S0047-6374(00)00118-4

